# Fisetin Attenuates Cellular Senescence Accumulation During Culture Expansion of Human Adipose-Derived Stem Cells

**DOI:** 10.1101/2022.11.15.516580

**Authors:** Michael Mullen, Alexander Goff, Jake Billings, Heidi Kloser, Charles Huard, John Mitchell, William Sealy Hambright, Sudheer Ravuri, Johnny Huard

## Abstract

Mesenchymal stem cells (MSCs) have long been viewed as a promising therapeutic for musculoskeletal repair. However, regulatory concerns including tumorgenicity, inconsistencies in preparation techniques, donor-to-donor variability, and the accumulation of senescence during culture expansion have hindered the clinical application of MSCs. Senescence, in particular, is a driving mechanism for MSC dysfunction with advancing age. Often characterized by increased reactive oxygen species, senescence-associated heterochromatin foci, inflammatory cytokine and chemokine secretion, and reduced proliferative capacity, senescence directly inhibits MSCs efficacy as a therapeutic for musculoskeletal regeneration and repair. Furthermore, autologous delivery of senescent MSCs can further induce disease and aging progression through the secretion of the senescence-associated secretory phenotype (SASP) and mitigate the regenerative potentetial of MSCs. To combat these issues, the use of senolytic agents to selectively clear senescent cell populations has become popular. However, their benefits to human MSCs during the culture expansion process have not yet been elucidated. To address this, analyzed markers of senescence during culturing of human primary adipose-derived stem cells (ADSCs), a population of fat-resident MSCs commonly used in regenerative medicine applications. Next, we used the senolytic agent fisetin to determine if we can reduce these markers of senescence within our culture-expanded ADSC populations. Our results indicate that ADSCs acquire common markers of cellular senescence including increased reactive oxygen species, senescence-associated β-galactosidase, and senescence-associated heterochromatin foci. Furthermore, we found that the senolytic agent fisetin works in a dose-dependent manner and selectively attenuates these markers of senescence while maintaining the differentiation potential of the expanded ADSCs.

**Signficance Statement:** The accumulation of dysfunctional, senescent cells throughout aging is not confined to specific tissues and cell types, but instead effects the whole body, including stem cells. Similarly, during culture expansion stem cells accumulate senescence while concurrently losing their regenerative potential. In this study, we found that fisetin (a well known senotherapeutic agent) can reduce the number of senescent cells during stem cell expansion. The current results indicate that fisetin may be used not only as a promising therapeutic to remove senescent cells in stem cell isolates from older individuals but also to reduce the accumulation of senescence during culture expansion.

## Introduction

In the 1960s and 70s, Dr. A.J. Friedenstein first identified spindle-shaped, plastic adherent cells from bone marrow (1). Later work would demonstrate both the osteogenic and chondrogenic potential of these cells (2, 3), and in 1991 this cell population would first be termed Mesenchymal Stem Cells (MSCs) by Dr. Arnold Caplan (4). Building on these seminal pieces of work, further evidence has demonstrated MSCs promise as a regenerative product across musculoskeletal tissues, nervous tissue, skin, and myocardium, among others (5). Recent evidence suggests that MSCs are a particularly promising therapeutic candidate due to their broad distribution in the body, diverse multilineage differentiation potential, and unique immunomodulatory characteristics (5, 6). Although both significant and exciting progress has been made in MSC-based therapeutic approaches, there are still several risks such as tumorigenicity and inconsistencies in preparation techniques and clinical outcomes that must be resolved to designate MSC-based therapies as reliable and effective treatment strategies to repair tissue after injury and disease. Therefore, improved and standardized techniques for MSC isolation, culture, and delivery are crucial to advancing our understanding of the mechanisms underlying the regenerative benefits of MSCs *in vivo*.

Originally discovered in bone marrow, MSCs have since been found in skeletal muscle (7), dental (8), umbilical cord (9), peripheral blood (10), and adipose tissues (11). However, bone marrow-derived MSCs (BMSCs), among others, are difficult to harvest and isolate, highlighting the need to identify novel, easily harvested repositories of MSCs (12). In this way, adipose tissue has evolved as a promising source of MSCs. Otherwise known as adipose-derived stem cells (ADSCs), MSCs from adipose tissue, exhibit similar multipotent potential as BMSCs but are significantly easier to harvest (12, 13) from subcutaneous fat. In healthy individuals, adipose tissue is systemically distributed, constitutes 10-30% of body weight, and lends a high cellular isolation yield per gram of harvested tissue (12, 14). Thus, ADSC isolation is both less invasive and more efficient than that of BMSCs, establishing ADSCs as an exciting biologic treatment option and regenerative therapy for age-related musculoskeletal conditions. The therapeutic potential of ADSCs for musculoskeletal conditions such as osteoarthritis (15), bone defects (16), and tendinopathy (17) have been investigated through a variety of pre clinical work and clinical trials. Initial results from these ADSC-based clinical trials are extremely promising; however, there are still limitations to their therapeutic potential *in vivo*.

To be considered a potential therapy for musculoskeletal repair, MSCs and ADSCs alike require processing beyond the current FDA guidelines for minimal manipulation and clinical administration (18). This is primarily due to the isolation and culture expansion required to create an administrable population of cells. Unfortunately, once removed from the body and cultured *in vitro*, previous research has shown that due to the *in vitro* culture conditions MSCs acquire new phenotypes including a modulated secretome, transcriptome, and surface epitope profile (19, 20). Similarly, due to differences in donor lifestyle, medical history, and age MSCs may not always elicit the same regenerative properties from donor to donor (21). These cell culture-acquired phenotypes and donor-dependent differences have been shown to result in a loss of proliferative and differentiation potential, as well as increased markers of senescence throughout the expansion process (22).

The accumulation of senescence *in vitro* was first observed in 1961 when Dr. Fredrick Hayflick observed human fibroblasts had a limited proliferative period during culture expansion (23). Biologically, senescence has several roles, including a protective mechanism to prevent the replication of damaged and/or mutated DNA (24). However, senescence is increasingly accepted to be a driving mechanism of aging where stressed and damaged cells enter a non-proliferative state (25). Senescent cell accumulation has also been shown to result in increased reactive oxygen species, senescence-associated β-galactosidase, senescence-associated heterochromatin foci, and a senescence-associated secretory phenotype (SASPs); which promote inflammation, degrade proteins, and leads to tissue dysfunction and further senescence induction to surrounding cells and tissues (22). Furthermore, the delivery of aged and/or senescent cells has been shown to diminish the maintenance, regeneration, and repair of musculoskeletal tissues *in vivo* (26, 27). Thus, the regenerative potential of MSCs is limited by issues with the conflicting goals of establishing a clinically effective MSC population via *in vitro* isolation and expansion while concurrently preventing harmful senescence accumulation.

The loss in the therapeutic potential of MSCs due to aging and culture expansion has given rise to the idea that senolytic agents may be used to rescue previously unusable donations by attenuating senescence and producing an enriched stem cell population. Moreover, numerous naturally occurring senolytic agents such as piperlongumine, dasatinib/quercetin, and fisetin have been found to exhibit senolytic activity by targeting anti-apoptotic pathways known to be upregulated in senescent cells (28-31). Concurrently, several reports have found significant improvement in healthspan indices *in vivo* following treatment with senolytic agents (32). Our group has demonstrated that fisetin, in particular, may be used to improve age-related cartilage and bone degeneration in a progeria mouse model (33). Additionally, these results further validate the use of fisetin as a senotherapeutic that extends health and lifespan in mice (34).

With this knowledge, we sought to investigate the effects of fisetin on senescence accumulation in MSCs using culture-expanded human adipose-derived stem cells (ADSCs). It was hypothesized that fisetin, a senolytic agent and antioxidant, can reduce senescent cell accumulation in ADSCs undergoing culture expansion while maintaining their regenerative potential. Ultimately, positive findings may highlight novel senotherapeutic strategies to improve the efficacy of stem cell-based therapies.

## Materials and Methods

### Cell Culture and Expansion

Banked adipose-derived stem cells (ADSCs) were purchased from CellTex Therapeutics (CellTex, catalog # N/A) and cultured in growth media containing DMEM-F:12 (Gibco, catalog # 11320033) with 10% Fetal Bovine Serum (FBS; Gibco, catalog # 10437028) and 1% penicillin/streptomycin (P/S; Gibco, catalog # 15140148). Donor ages ranged from 10 – 80 years of age. All cells were maintained in tissue culture-treated T75 flasks (Corning, catalog # 10-126-11) at 37°C in humidified air containing 5% CO_2_. All cells were received at passage 1 and subsequently passaged up to 18 times. At 70-90% confluency, the cells were passaged and reseeded at a 1:4 dilution.

### Determination of Cell Number and Proliferation

Cell number was determined by removing the cells using TrypLE (Gibco, catalog # 12605028) and conducting direct cell counts using the Countess II automated cell counter. The metabolic activity of the cells was quantified by treating cells with the PrestoBlue Cell Viability Reagent (Invitrogen, catalog # A13262) for two hours and measuring the optical density at 570 nm per manufacturer’s protocol. The growth rate was determined using the following equation:

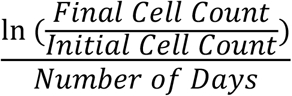

### Reactive Oxygen Species and C_12_FDG Quantification

Cells positive for reactive oxygen species and senescence-associated β-galactosidase were quantified using flow cytometry. Senescence-associated β-galactosidase was tested using the C_12_FDG substrate which becomes fluorescent when hydrolyzed by SA-β-galactosidase. ADSCs were tested at passages 4, 6, and 8 after being plated in 12-well plates (Genesee, catalog # 25106) in triplicate at a density of 40,000 cells/well. The cells were treated with 100nM of bafilomycin A1 (Cell Signaling, catalog # 54645) for 1 hour at 37°C and 5% CO_2_. Next, 33µM C_12_FDG (Thermo Fisher, catalog # D2893) was added and incubated for 2 hours. After 1.5 hours the CellROX reagent (Thermo Fisher, catalog # C10422) was also added for the remaining thirty minutes. The cells were then washed with PBS and collected from the plate using the TrypLE reagent. Using the Guava EasyCyte flow cytometer, cells positive for C_12_FDG and reactive oxygen species were quantified using unstained cells to set gate thresholds for fluorescence.

### Senescence Associated Heterochromatin Foci Staining

Cells were plated into chamber slides for senescence-associated heterochromatin foci (SAHF) staining at a density of 5,000 cells/cm^2^. Cells were seeded in triplicate for each group and incubated to adhere overnight. The following day, cells were fixed with cold 4% paraformaldehyde (Alfa Aeser, Catalog # J19943K2) for 15 minutes, and washed three more times using PBS. The cells were then blocked using 10% donkey serum (DS; Jackson Immuno, catalog # 017000121) and 0.3% Triton X-100 (Fisher, catalog # BP151-500) in PBS for 1 hour followed by overnight incubation at 4°C with the γ-H2AX antibody (Millipore Sigma, catalog # 05636I) using a 1:150 dilution and the H3K9me3 antibody (Millipore Sigma, catalog # 07442) using a 1:250 dilution. Both antibodies were added in dilution buffer containing 1X PBS with 1% DS, 1% bovine serum albumin (BSA; Sigma, catalog # A9647-100G), and 0.3% Triton X-100. Following antibody conjugation, the cells were washed three times with wash buffer containing 1X PBS with 0.1% BSA. The cells were then incubated with 1:400 diluted Alexa Flour 488 and 594 (Invitrogen, catalog # A21206 and A21203, respectively) secondary antibodies for 1 hour at room temperature. The cells were washed three more times and incubated with 1 µg/ml of DAPI (Sigma, catalog # D9542-10MG) for 10 minutes before being washed with PBS. Five images were taken per well and three wells per group were imaged using the Nikon Eclipse Ni-U microscope. Cells positive for the γ-H2AX and H3K9me3 antibodies were quantified using the ImageJ software.

### Fisetin Treatment

Cells were seeded in triplicate for each group using previously described culture methods and seeding densities. After the cells adhered to the flask, the media was changed. For the untreated group, normal ADSC growth media was used (DMEM:F-12 with 10% FBS and 1% penicillin/streptomycin), and for the treated groups 25µM, 50µM, or 100µM of fisetin (Selleckchem, catalog # S2298) was added. Cells were treated for 24-hours, after which the wells were washed and normal growth media was added.

### Adipogenic and Osteogenic Differentiation

ADSC differentiation was carried out using the Lonza Adipogenesis and Osteogenesis kits (Lonza, catalog # PT3004 and PT3002, respectively) according to the manufacturer’s instructions. 20,000 cells were seeded into each well of a 24-well plate (Genesee, catalog # 25-107). Cells were grown to confluency using normal growth media. Once all wells reached 90-100% confluency the media was changed to differentiation media. For adipogenic differentiation, the Lonza Adipogenic Induction Media was added for 3 days followed by 2 days with the Lonza Adipogenic Maintenance Media. This process was repeated for a total of 15 days. For osteogenic differentiation, fresh Lonza Osteogenic Induction Media was added every 3 days for a total of 15 days.

### RNA Isolation, Reverse Transcription, and Quantitative PCR

Total RNA was extracted using the Trizol reagent (Invitrogen, catalog # 15596026) according to the manufacturer’s instructions. Following RNA isolation, reverse transcription was performed using the qScrpit cDNA synthesis kit (VWR, catalog # 95048-100) according to the manufacturer’s instructions. Quantitative real-time PCR was performed using the PerfecCTa SYBR Green FastMix (VWR, catalog # 101414-280) on the Step-One Plus Real-Time PCR system (Applied Biosystems, catalog # 4376598). All results were normalized to the Gapdh reference gene. Primer sequences were as follows: Gapdh F: GGAGCGAGATCCCTCCAAAAT R: GGCTGTTGTCATACTTCTCATGG, PPARγ F: TACTGTCGGTTTCAGAAATGCC R: GTCAGCGGACTCTGGATTCAG, C/EBPα F: ACTCCAGGGGTGAACGGAAT R: CATGGGCGAACTCTTTTTGCT, FABP4 F: ACTGGGCCAGGAATTTGACG R: CTCGTGGAAGTGACGCCTT, Col1a F: AGGGCTCCAACGAGATCGAGATCCG R: TACAGGAAGCAGACAGGGCCAACGTCG, ALP F: GACCTCCTCGGAAGACACTC R: TGAAGGGCTTCTTGTCTGTG, OC F: GGCGCTACCTGTATCAATGG R: GTGGTCAGCCAACTCGTCA.

### Statistics

All statistical comparisons and figures were generated using the PRISM 9 analysis software. T-tests were completed for all comparisons between two groups. One-way ANOVAs were completed to test for significant differences between three or more groups, and two-way ANOVAs were completed to compare multiple groups over time. Tukey’s honestly significant difference (HSD) post-hoc testing was completed in both scenarios. Values with a p < 0.05 were considered to be statistically significant.

## Results

### Adipose-Derived Stem Cells Accumulate Markers of Cellular Senescence During Culture Expansion

Following extended culture expansion of banked human Adipose-Derived Stem Cells (ADSCs) one-way ANOVA results found that the growth rate significantly decreased over time (**Figure 1A**). This was particularly evident after passage 6 when the growth rate significantly declined when compared to passage 8 (p < 0.05). This decrease was also found to be significantly different between passage 4 and passage 8 (p < 0.01). Concurrent with the changes in growth rate, one-way ANOVA results found that the number of cells positive for reactive oxygen species (ROS) was significantly increased with passaging (**Figure 1B**). Between passage 4 and 6 a 4.4% ROS increase was observed (p < 0.0001) followed by a 5.7% increase when compared to passage 8 cells (p = 0.0008). Furthermore, senescence-associated marker β-galactosidase was upregulated in ADSCs after culture expansion (**Figure 1C**). Here, we observed a significant increase between passage 4 and 6 (Δ = 15.48%, p = 0.018), 4 and 8 (Δ = 43.46%, p = 0.0032), and 6 and 8 (Δ = 27.98%, p < 0.0001).

**Figure 1:**
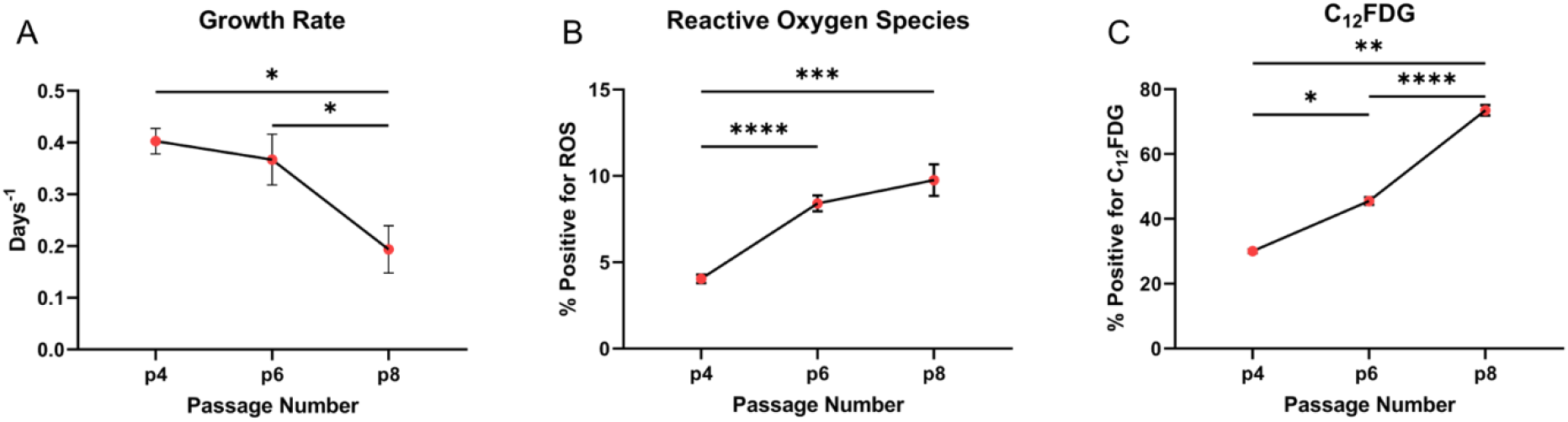
Adipose Derived Stem Cells Lose Function and Acquire Markers of Senescence During Culture Expansion. A) Growth rate significantly decreases with culture expansion. B) Cells positive for reactive oxygen species increase with culture expansion. C) Cells positive for the C_12_FDG marker increase with culture expansion. (* = p < 0.05, ** = p < 0.01, *** = p < 0.001, **** = p < 0.0001).

Senescence-associated heterochromatin foci (SAHF) are a common feature during senescence progression, thus we next investigated the changes observed between low and high ADSCs passage (**Figure 2**). When comparing passage 4 and passage 18 ADSCs, we observed that passage 18 presented significantly more cells positive for H3K9me3 than ADSCs at passage 4 (Δ = 31.17%, p = 0.014, **Figure 2A**). Similarly, within the cells positive for H3K9me3, significantly more foci were found within passage 18’s nuclei when compared to passage 4 (Δ = 6.52 foci/nuclei, p = 0.014, **Figure 2B**). When investigating the changes associated with SAHF γ-H2AX an increase in γ-H2AX positive cells at passage 18 cells was observed, albeit not significantly (Δ = 16.00%, p = 0.078, **Figure 2C**). However, within the nuclei positive for γ-H2AX, significantly more foci were observed in ADSCs at passage 18 (Δ = 5.45 foci/nuclei, p = 0.0043, Figure 2D) when compared to passage 4 ADSCs.

**Figure 2:**
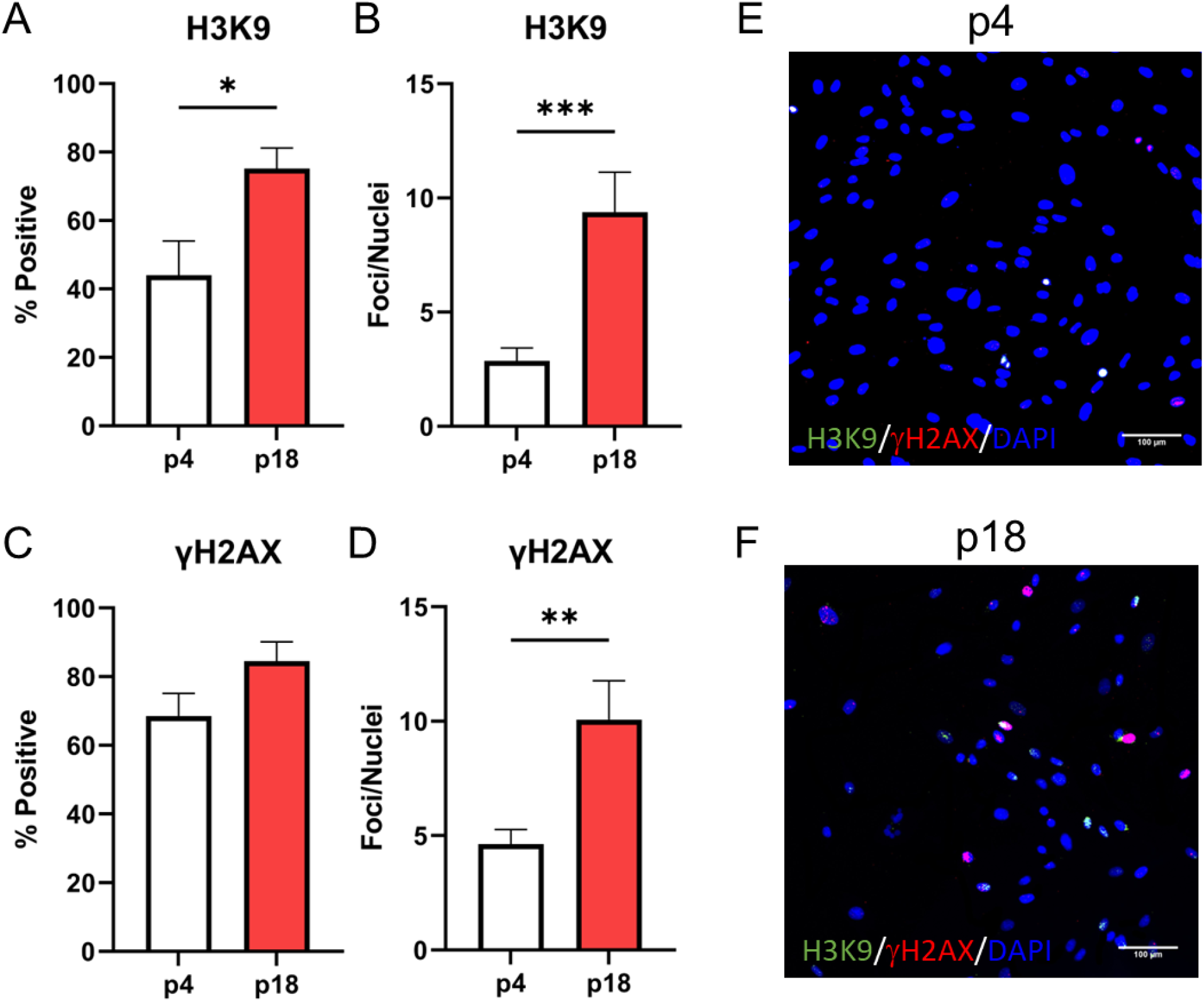
Senescence Associated Heterochromatin Foci Increase in High Passage ADSC. A-B) Cells positive for H3K9 and the number of H3K9 foci per nuclei are upregulated with passaging. C-D) Cells positive for γ-H2AX and the number of γ-H2AX foci per nuclei are upregulated with passaging. E-F) Representative immunofluorescence images of low and high passage senescence associated heterochromatin foci. (* = p < 0.05, ** = p < 0.01, *** = p < 0.001)

### Fisetin Attenuates Cellular Senescence in a Dose-Dependent Manner in ADSCs

After observing the accumulation of senescent cells during culture expansion on ADSCs, we next looked to determine the potential use of fisetin in eliminating senescent cells during culture expansion. To determine this, we used the PrestoBlue Cell Viability Reagent to determine the relative number of cells removed following treatment with increasing concentrations of fisetin (**Figure 3A**). Following a 24-hour treatment and 24-hour recovery, we found a significant decrease in cell viability with increasing concentrations of fisetin. To determine if the reduction in cell number corresponded with a reduction in potential senescent cells, we used the C_12_FDG substrate to quantify the number of cells presenting senescence-associated β-galactosidase (**Figure 3B**). These results correlated with the reduction in cell viability indicating that fisetin selectively targeted senescent cells. From these results, we found that the 50µM fisetin concentration would be optimal as it removed 43.7% of senescent cells while cell viability was only reduced by 17.5% compared to untreated cells (**Figure 3A-C**). Comparing this to the 100µM concentration, no significant difference was found in the reduction of cells positive for C_12_FDG when compared to the 50µM concentration (Δ = 4.7%, p = 0.1258). However, there was a significant difference in the total cell viability when comparing the 50µM and 100µM fisetin concentrations (Δ = 29.9%, p < 0.0001), leading us to believe that 100µM of fisetin may have also eliminated non-senescent proliferating cells.

**Figure 3:**
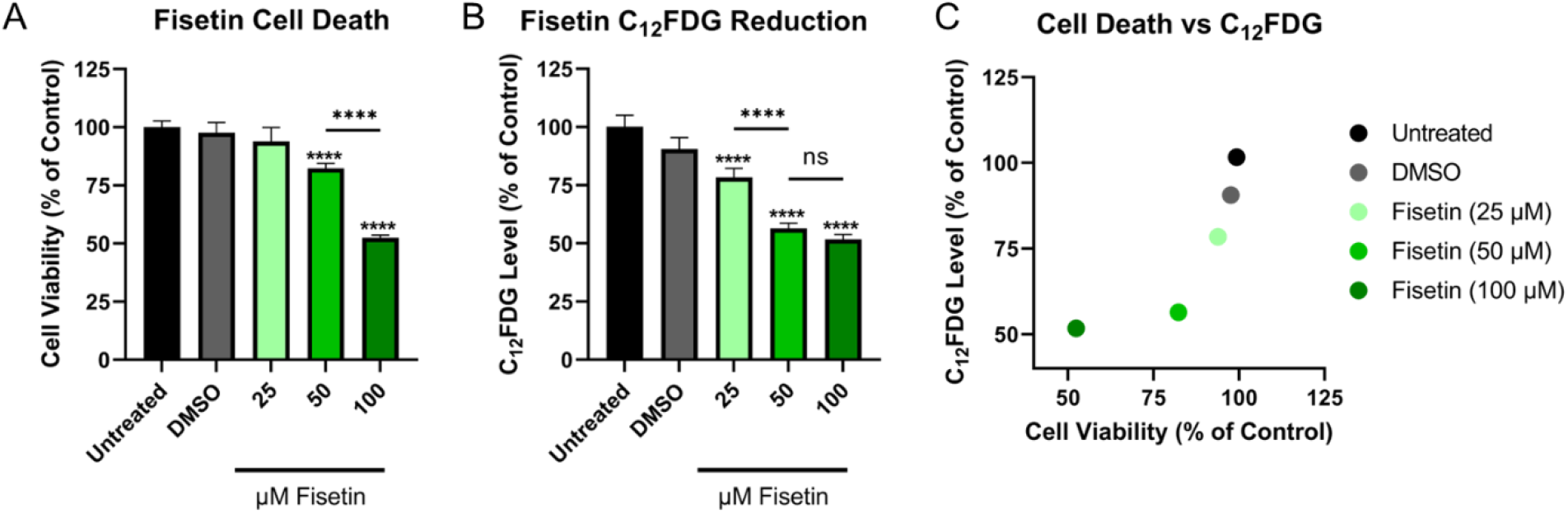
Effect of Flavonoids on ADSC Cell Death and Senescence. A) Fisetin induces cell death with increasing concentration when compared to untreated ADSCs. B) Fisetin reduces C_12_FDG positive ADSC populations with increasing concentration. C) Cell viability is plotted against the corresponding reduction in C_12_FDG following fisetin treatment. (**** = p < 0.0001)

Based on the results found in figure 3 we used 50µM of fisetin to determine if we could attenuate other markers of senescence at both low and high passage from multiple ADSC samples. Here we found that again 50µM of fisetin reduced the number of cells with elevated reactive oxygen species by 73.8% in passage 4 cells and 78.4% in passage 10 cells (p < 0.0001, **Figure 4B**). Similarly, the number of cells positive for β-galactosidase, as determined by C_12_FDG quantification, was reduced by 40.9% in passage 4 cells and 49.0% in passage 10 cells (p < 0.0001, **Figures 4F**).

**Figure 4:**
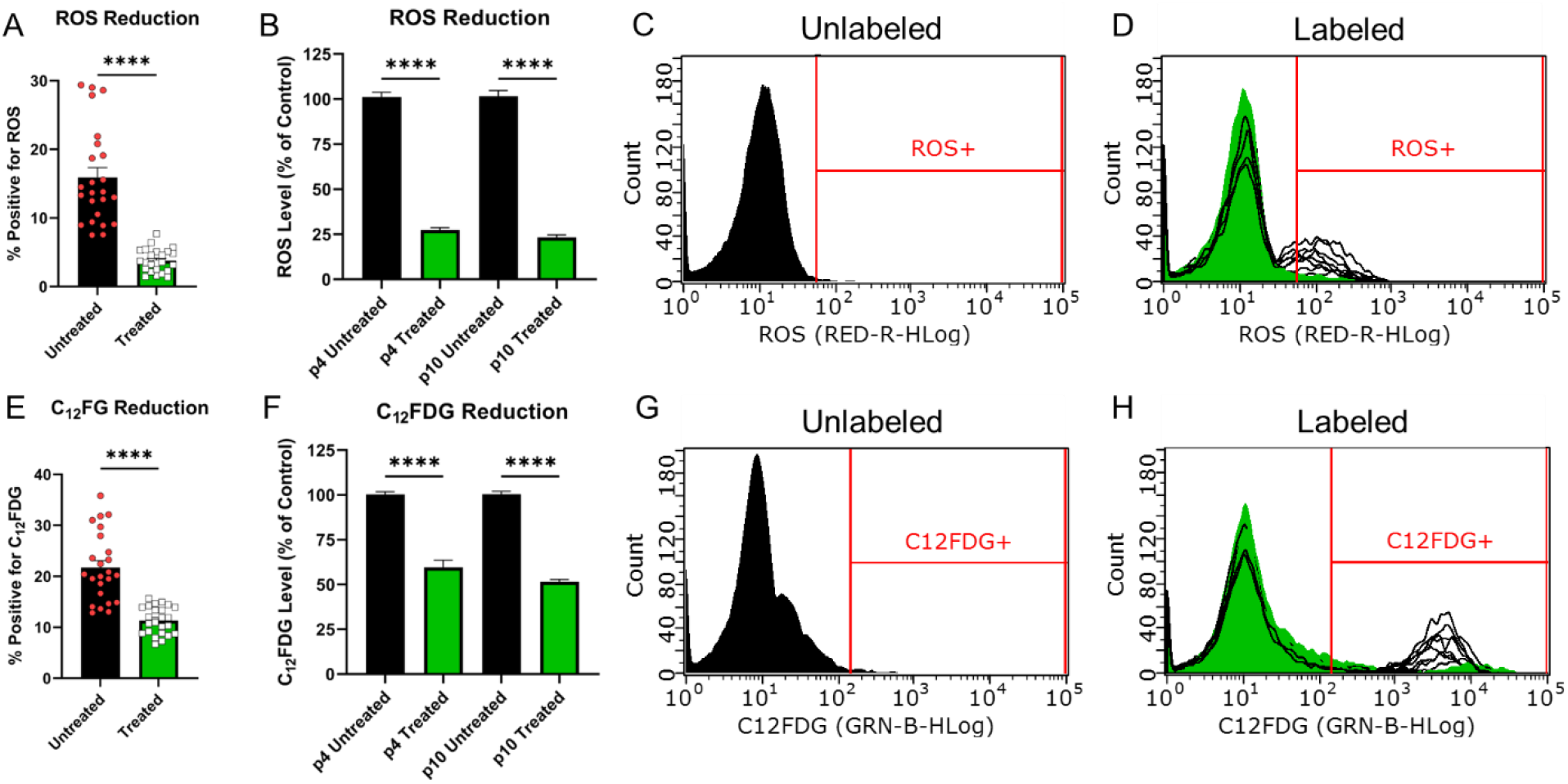
Fisetin Treatment Reduces Reactive Oxygen Species and C_12_FDG. Cells were treated with 50µM of Fisetin and tested for ROS and C_12_FDG using flow cytometry. A) ROS positive untreated ADSCs compared to treated. B) ROS reduction relative to untreated at passage 4 (p4) and passage 10 (p10). E) C_12_FDG positive untreated ADSCCs compared to treated. F) C_12_FDG reduction relative to untreated at p4 and p10. Black peaks in histograms represent unlabeled cells while white and green peaks represent untreated and treated, respectively. (**** = p < 0.0001)

Following ROS and C_12_FDG quantification, we determined the number of ADSCs positive for H3K9me3 and γ-H2AX SAHF following treatment with 50µM of fisetin. Here, we again observed a significant decrease with 58.2% and 57.3% fewer cells presenting H3K9me3 SAHF at passages 4 and 18, respectively (p < 0.001, **Figure 5A**). Furthermore, within the cells positive for H3K9me3 foci following treatment with fisetin, the number of foci present was significantly reduced in passages 4 and 18 (Δ = 68.0% p = 0.0086 and Δ = 73.7% p = 0.0037, respectively, **Figure 5B**). These results were conserved when quantifying the number of γ-H2AX foci. Here we observed 36.3% (p = 0.0004) and 37.6% (p = 0.0002) fewer nuclei positive for γ-H2AX foci when compared to untreated controls at passage 4 and 18 respectively (**Figure 5C**). Again, among the nuclei presenting γ-H2AX foci a significant reduction in the number of foci/nuclei was observed in both passages 4 and 18 ADSCs after treatment (Δ = 55.5% p = 0.0069 and Δ = 59.4% p = 0.0033, respectively, **Figure 5D**).

**Figure 5:**
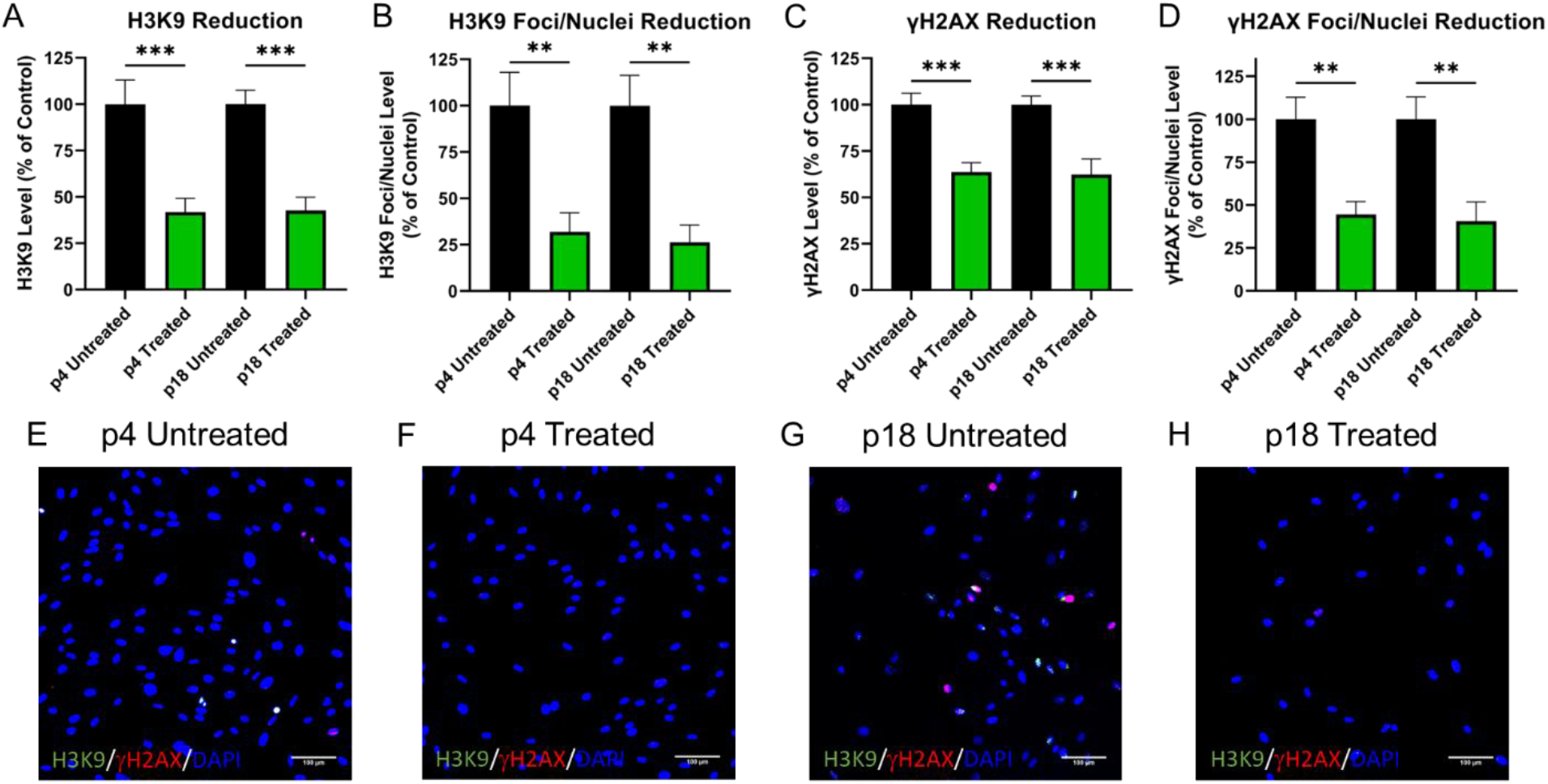
Fisetin Treatment Reduces Senescence Associated Heterochromatin Foci. Cells were treated with 50µM of fisetin and tested for H3K9 and γ-H2AX. A-B) H3K9 positive cells and H3K9 foci/nuclei are reduced following fisetin treatment. C-D) γ-H2AX positive cells and γ-H2AX foci/nuclei are reduced following fisetin treatment. E-H) Representative immunofluorescence images of passage 4 and 18 treated and untreated cells. H3K9 = Green, γ-H2AX = Red, and Nuclei = Blue. (** = p < 0.01, *** = p < 0.001)

### Fisetin Treatment Maintains Adipose-Derived Stem Cell Adipogenic and Osteogenic Differentiation Capacity

To determine the effects of fisetin on the multipotent differentiation of ADSCs, we performed adipogenic and osteogenic differentiation following treatment with 50µM of fisetin on low passage (p4) cells. We found that both adipogenesis and osteogenesis were unaffected by fisetin treatment. Adipogenic transcripts PPARγ, C/EBPα, and FABP4 were all not significantly different following 15 days of adipogenic differentiation when compared to untreated cells (p = 0.3803, 0.2547, and 0.6967, respectively, **Figure 6A-C**). Similarly, oil red o staining for lipogenesis showed no significant change in lipid vesicle production between treated and untreated cells (p = 0.8330, **Figure 6D-E**). The osteogenic potential was also not altered by fisetin treatment as determined by qPCR and alizarin red staining (**Figure 7**). Following 15 days of differentiation, collagen type 1 (Col1a), osteocalcin (OC), and alkaline phosphatase (ALP) were all similarly expressed between treated and untreated cells (p = 0.3568, 0.3583, 0.5906, respectively, **Figure 7A-C**). Alizarin red staining was also unaltered between treated and untreated cells (p = 0.0755, **Figure 7D-F**).

**Figure 6:**
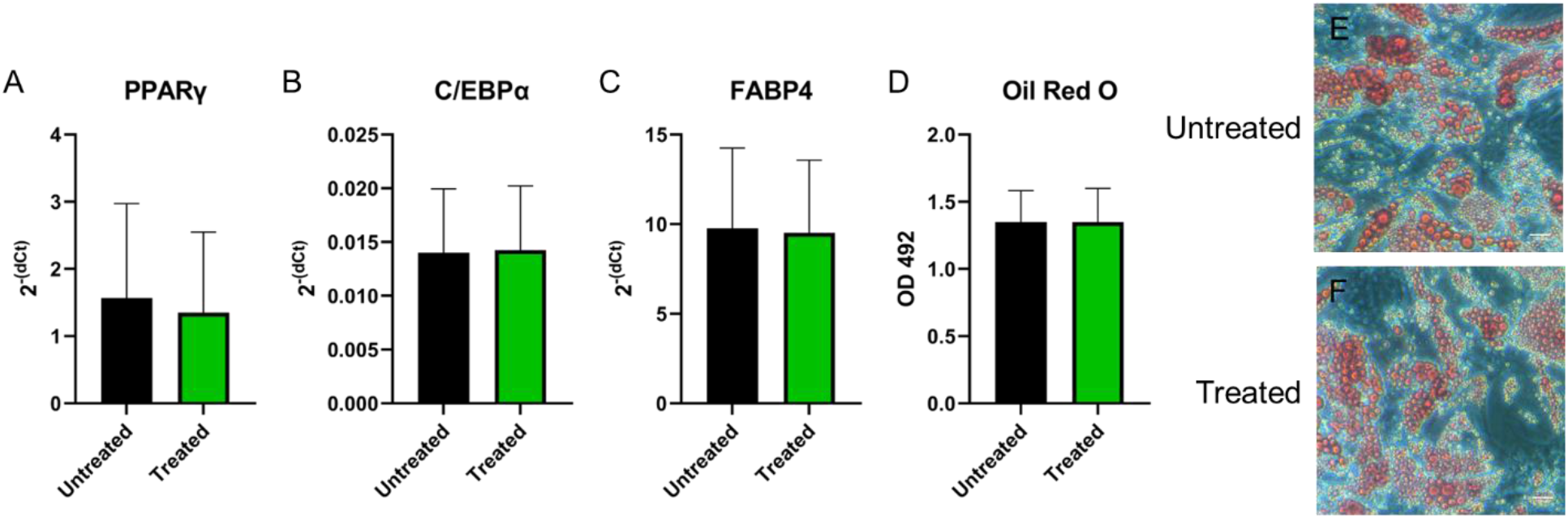
Fisetin Treatment Maintains Adipogenic Potential. A-C) Adipogenic markers PPARγ, C/EBPα and FABP4 are not significantly affected by fisetin treatment. D) Lipid production is not significantly altered after 50µM Fisetin treatment. E-F) Representative Oil Red O images following 15 days of differentiation.

**Figure 7:**
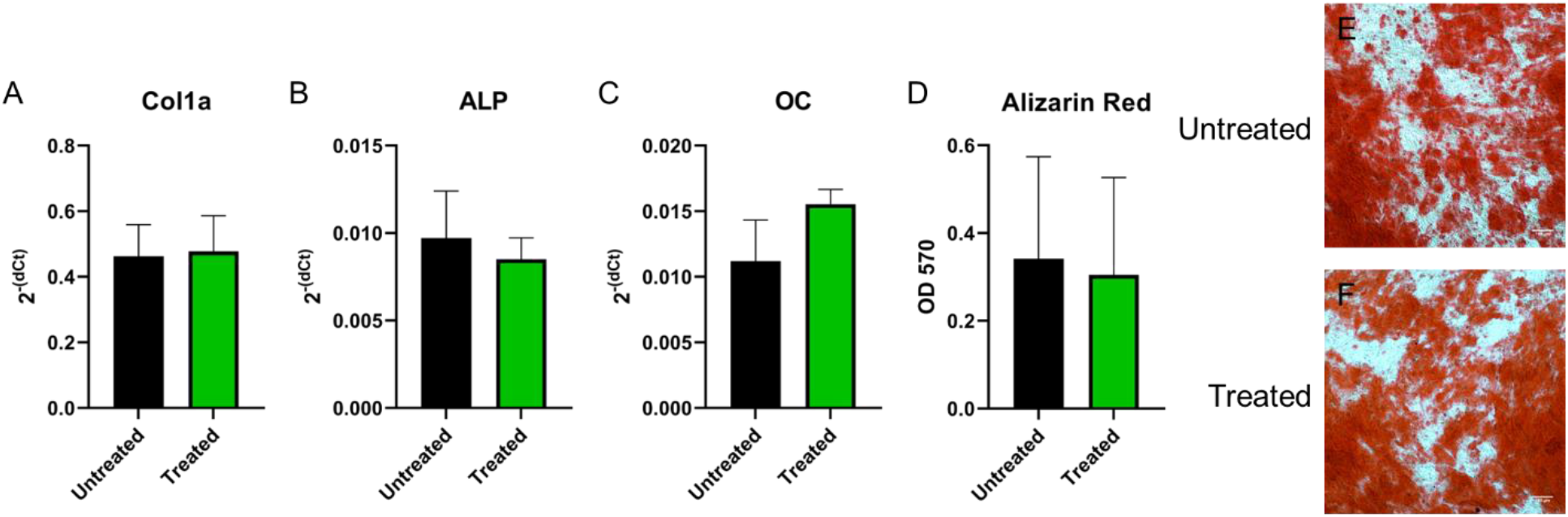
Fisetin Treatment Maintains Osteogenic Potential. A-C) Osteogenic markers Col1a, ALP, and OC are not significantly affected by fisetin treatment. D) Calcium production is not significantly altered after 50µM fisetin treatment. E-F) Representative alizarin red images following 15 days of differentiation.

## Discussion

The regenerative potential of human mesenchymal stem cells shows promise in the field of musculoskeletal tissue regeneration with several reports demonstrating the potential of MSCs to promote musculoskeletal tissue healing and repair (35, 36). However, for many regenerative medicine therapies, MSCs are often cultivated for several passages *in vitro* to achieve the number of cells required for transplantation (37). It has been demonstrated that culture expansion affects the regenerative potential of adult stem cells (38, 39). Cellular senescence is thought to be a fundamental driver of aging and a major contributor to age-associated decline and loss of physiological reserve (25). Senescent cell accumulation is not only a fundamental property of aging, but also promotes several age-related pathologies and can affect a person’s ability to withstand stress, recuperate from injury, and operate at peak mental capacity (24, 25, 40). There are, however, drugs and supplements (“senolytics”) that can selectively target senescent cells, reducing their harmful effects, even in older individuals (29, 31). Here, we aimed at characterizing cellular senescence during the expansion of human adipose-derived stem cells (ADSCs) and tested whether treatment with the senolytic fisetin could reduce the accumulation of senescence during culture expansion.

To assess the impacts of culture expansion on senescence accumulation in MSCs and ADSCs alike, we investigated common markers associated with senescence at different passages of ADSCs. Due to the elusive pathology of senescence both *in vitro* and *in vivo*, it was known that not all markers referenced in the literature may be present. In previous studies it was shown that the p53 pathway triggers the cellular senescence process in MSCs (41), thereby resulting in the accumulation of non-proliferative senescent MSCs (22, 42, 43). Furthermore, the expression of reactive oxygen species, senescence-associated β-galactosidase, DNA damage, nuclear foci such as γ-H2AX, and a complex of growth factors i.e., proteases and cytokines (which are collectively called the senescence-associated secretory phenotypes (SASPs)) have shown to be hallmark makers of senescence (44). In the present study, we identified several of these canonical markers of senescence, including reactive oxygen species, senescence-associated β-galactosidase, and senescence-associated heterochromatin foci, to be associated with human ADSCs undergoing extended culture expansion.

Since MSCs attain replicative senescence during *in vitro* cultural expansion it is of critical importance to address these challenges during MSC expansions. There is a need to standardize the regulatory criteria as cellular senescence can impair the ability of MSCs to suppress inflammation and/or reduce their therapeutic efficacy in tissue repair. The study by Wagner et al. showed that culture expansion of MSCs accumulated gradual changes in the global gene and miRNA expression (43). Furthermore, the study findings concluded that the senescent-associated changes that occurred in gene and protein expressions were not only associated with culture expansion but were also observed at the beginning of *in vitro* culture. Hence, identifying, characterizing, and/or eliminating senescent cells in early MSC cultures can be an important step to ensure the best MSC product for cell-based therapy.

Targeting and safely eliminating senescent cells with senolytic agents is now widely studied as cellular senescence is emerging as the conduit to a multitude of age related disease conditions such as osteoarthritis (27), Alzheimer’s disease (45), obesity (46), pulmonary fibrosis (47), and chronic kidney disease (48). Senolytics are a class of drugs that selectively clear or eliminate senescence by inducing apoptosis in senescent cells. In particular, dasatinib, quercetin, fisetin, and navitoclax have all been shown to eliminate senescence in many preclinical models by delaying or alleviating frailty, cancer, and cardiovascular disease (34, 49, 50).

Of the numerous senolytic agents discovered to date, fisetin is of particular interest due to its established antioxidant, anticarcinogenic, anti-inflammatory, and apoptotic qualities (51). Fisetin is a bioactive flavonol molecule found in fruits and vegetables such as cucumber, apple, grape, and onion, with the highest concentration being found in strawberries (51). Aged C57BL/6 mice treated orally at 22–24 months with 100 mg/kg fisetin for 5 days showed a reduction in senescent cells in white adipose tissue (34). Additionally, fisetin treatment of mice, at 85 weeks of age, significantly prolonged the lifespan of these mice by an additional 3 months. This research has given rise to the Alleviation by Fisetin of Frailty, Inflammation and Related Measures in Older Adults (AFFIRM-LITE) clinical trial. Currently, in the recruiting phase, the trial hopes to recruit 40 participants between the ages of 70–90 to take an oral 2-day dose of placebo or fisetin at 20 mg/kg/day with analysis focusing on markers of frailty, inflammation, insulin resistance, and bone metabolism (52). Previously, it has been shown that fisetin can act as a senolytic both *ex vivo* and *in vivo*. Currently, our study sheds light on the use of fisetin *in vitro* to reduce senescence accumulation during the culture expansion of MSCs.

In conclusion, typically, freshly isolated stem cell numbers are limited and it is necessary to expand their populations *in vitro* before clinical use. To examine the characteristics and safety of long-term cultured MSCs, the current study evaluated the effects of long-term culture on MSC senescence accumulation. These results indicate that MSCs undergo replicative senescence during long-term culture *in vitro*, as demonstrated by decreases in growth rate with concurrent increases in reactive oxygen species, senescence-associated β-galactosidase, and senescence-associated heterochromatin foci. However, in all cells, baseline levels of all markers were noted, indicating that the quality of MSC preparations should be carefully assessed before clinical application. To mitigate these markers, this study also demonstrated the use of single treatments of the senolytic agent fisetin in removing senescent cell populations in both low and high passage preparations without affecting their multipotent differentiation potential; providing an important insight into enriching freshly isolated orthobiologics, such as stem cells or bone marrow concentrate, by treating the biologic product with fisetin before banking or administration to the patient. While the current results are promising for future improvements to stem cell therapies, further research will be necessary to understand the mechanisms that regulate the replicative senescence of stem cells and how to ensure MSCs remain in a senescent-free state during culture expansion.

## Acknowledgments

We would like to thank Suzanne Page for her administrative support.

## Conflicts of Interest

Michael Mullen, Alexander Goff, Jake Billings, Charles Huard, John Mitchell, Heidi Kloser, Dr. William Seally Hambright, and Dr. Sudheer Ravuri have no conflicts of interest. Dr. Johnny Huard discloses an unpaid position on the leadership board of the Orthopaedic Research Society (ORS). JH discloses royalties from Cook Myosite, Inc. All authors are or were paid employees of the non-profit Steadman Philippon Research Institute (SPRI). SPRI exercises special care to identify any financial interests or relationships related to research conducted here. During the past calendar year, SPRI has received grant funding or in-kind donations from Arthrex, DJO, MLB, Ossur, Siemens, Smith & Nephew, XTRE, and philanthropy. These funding sources provided no support for the work presented in this manuscript unless otherwise noted.

## Data Availability Statement

The data underlying this article will be shared on reasonable request to the corresponding author.

## Notes

**Source of Funding** This study was generously supported through a philanthropic donation from the Borgen family and the Linda and Mitch Hart Foundation.

## References

1. Friedenstein AJ. Osteogenic Stem Cells in Bone Marrow. Bone and Mineral Research. 1990:243–72.

2. Ashton BA, Allen TD, Howlett CR, Eaglesom CC, Hattori A, Owen M. Formation of bone and cartilage by marrow stromal cells in diffusion chambers in vivo. Clinical orthopaedics and related research. 1980(151):294–307.

3. Owen M, Friedenstein AJ. Stromal Stem Cells: Marrow-Derived Osteogenic Precursors. 2007. p. 42–60.

4. Caplan AI. Mesenchymal stem cells. Journal of orthopaedic research : official publication of the Orthopaedic Research Society. 1991;9(9):641–50.

5. Andrzejewska A, Lukomska B, Janowski M. Concise Review: Mesenchymal Stem Cells: From Roots to Boost. STEM CELLS. 2019;37(37):855–64.

6. Parekkadan B, Milwid JM. Mesenchymal Stem Cells as Therapeutics. Annual Review of Biomedical Engineering. 2010;12(12).

7. Jankowski RJ, Deasy BM, Huard J. Muscle-derived stem cells. Gene Therapy. 2002;9(9):642–7.

8. Gronthos S, Mankani M, Brahim J, Robey PG, Shi S. Postnatal human dental pulp stem cells (DPSCs) in vitro and invivo. Proceedings of the National Academy of Sciences. 2000;97(97):13625–30.

9. Erices A, Conget P, Minguell JJ. Mesenchymal progenitor cells in human umbilical cord blood. British Journal of Haematology. 2000;109(109):235–42.

10. Roufosse CA, Direkze NC, Otto WR, Wright NA. Circulating mesenchymal stem cells. The International Journal of Biochemistry & Cell Biology. 2004;36(36):585–97.

11. Zuk PA, Zhu M, Mizuno H, Huang J, Futrell JW, Katz AJ, et al. Multilineage Cells from Human Adipose Tissue: Implications for Cell-Based Therapies. Tissue Engineering. 2001;7(7):211–28.

12. Usuelli FG, D’Ambrosi R, Maccario C, Indino C, Manzi L, Maffulli N. Adipose-derived stem cells in orthopaedic pathologies. British Medical Bulletin. 2017:1–24.

13. De Ugarte DA, Morizono K, Elbarbary A, Alfonso Z, Zuk PA, Zhu M, et al. Comparison of Multi-Lineage Cells from Human Adipose Tissue and Bone Marrow. Cells Tissues Organs. 2003;174(174):101–9.

14. Fraser JK, Wulur I, Alfonso Z, Hedrick MH. Fat tissue: an underappreciated source of stem cells for biotechnology. Trends in Biotechnology. 2006;24(24):150–4.

15. Kim YS, Choi YJ, Suh DS, Heo DB, Kim YI, Ryu J-S, et al. Mesenchymal Stem Cell Implantation in Osteoarthritic Knees. The American Journal of Sports Medicine. 2015;43(43):176–85.

16. Dufrane D, Docquier P-L, Delloye C, Poirel HA, André W, Aouassar N. Scaffold-free Three-dimensional Graft From Autologous Adipose-derived Stem Cells for Large Bone Defect Reconstruction. Medicine. 2015;94(94):e2220–e.

17. Usuelli FG, Grassi M, Maccario C, Vigano’ M, Lanfranchi L, Alfieri Montrasio U, et al. Intratendinous adipose-derived stromal vascular fraction (SVF) injection provides a safe, efficacious treatment for Achilles tendinopathy: results of a randomized controlled clinical trial at a 6-month follow-up. Knee Surgery, Sports Traumatology, Arthroscopy. 2018;26(26):2000–10.

18. Regulatory Considerations for Human Cells, Tissues, and Cellular and Tissue-Based Products: Minimal Manipulation and Homologous Use. 2020.

19. Truong NC, Bui KH-T, Van Pham P. Characterization of Senescence of Human Adipose-Derived Stem Cells After Long-Term Expansion. 2018.

20. DiGirolamo CM, Stokes D, Colter D, Phinney DG, Class R, Prockop DJ. Propagation and senescence of human marrow stromal cells in culture: a simple colony-forming assay identifies samples with the greatest potential to propagate and differentiate. British Journal of Haematology. 1999;107(107):275–81.

21. Baker N, Boyette LB, Tuan RS. Characterization of bone marrow-derived mesenchymal stem cells in aging. Bone. 2015;70:37–47.

22. Liu J, Ding Y, Liu Z, Liang X. Senescence in Mesenchymal Stem Cells: Functional Alterations, Molecular Mechanisms, and Rejuvenation Strategies. Frontiers in Cell and Developmental Biology. 2020;8.

23. Hayflick L, Moorhead PS. The serial cultivation of human diploid cell strains. Experimental Cell Research. 1961;25(25):585–621.

24. Kowald A, Passos JF, Kirkwood TBL. On the evolution of cellular senescence. Aging Cell. 2020;19(19).

25. LeBrasseur NK, Tchkonia T, Kirkland JL. Cellular Senescence and the Biology of Aging, Disease, and Frailty. p. 11–8.

26. Lavasani M, Robinson AR, Lu A, Song M, Feduska JM, Ahani B, et al. Muscle-derived stem/progenitor cell dysfunction limits healthspan and lifespan in a murine progeria model. Nature Communications. 2012;3(3):608-.

27. Xu M, Bradley EW, Weivoda MM, Hwang SM, Pirtskhalava T, Decklever T, et al. Transplanted Senescent Cells Induce an Osteoarthritis-Like Condition in Mice. The Journals of Gerontology Series A: Biological Sciences and Medical Sciences. 2016:glw154–glw.

28. Wang Y, Chang J, Liu X, Zhang X, Zhang S, Zhang X, et al. Discovery of piperlongumine as a potential novel lead for the development of senolytic agents. Aging. 2016;8(8):2915–26.

29. Fuhrmann-Stroissnigg H, Ling YY, Zhao J, McGowan SJ, Zhu Y, Brooks RW, et al. Identification of HSP90 inhibitors as a novel class of senolytics. Nature Communications. 2017;8(8):422-.

30. Zhu Y, Doornebal EJ, Pirtskhalava T, Giorgadze N, Wentworth M, Fuhrmann-Stroissnigg H, et al. New agents that target senescent cells: the flavone, fisetin, and the BCL-XL inhibitors, A1331852 and A1155463. Aging. 2017;9(9):955–63.

31. Kirkland JL, Tchkonia T. Senolytic drugs: from discovery to translation. Journal of Internal Medicine. 2020;288(288):518–36.

32. Xu M, Pirtskhalava T, Farr JN, Weigand BM, Palmer AK, Weivoda MM, et al. Senolytics improve physical function and increase lifespan in old age. Nature Medicine. 2018;24(24):1246–56.

33. Hambright S, Kawakami Y, Mu X, Gao X, Lu A, Kirkland J, et al. The senolytic drug fisetin mitigates age-related bone density loss in the progeroid mouse model Zmpste24-/-. The FASEB Journal. 2020;34(S1):1-.

34. Yousefzadeh MJ, Zhu Y, McGowan SJ, Angelini L, Fuhrmann-Stroissnigg H, Xu M, et al. Fisetin is a senotherapeutic that extends health and lifespan. EBioMedicine. 2018;36:18–28.

35. Berebichez-Fridman R, Gómez-García R, Granados-Montiel J, Berebichez-Fastlicht E, Olivos-Meza A, Granados J, et al. The Holy Grail of Orthopedic Surgery: Mesenchymal Stem Cells-Their Current Uses and Potential Applications. Stem cells international. 2017;2017:2638305-.

36. Wei C-C, Lin AB, Hung S-C. Mesenchymal Stem Cells in Regenerative Medicine for Musculoskeletal Diseases: Bench, Bedside, and Industry. Cell Transplantation. 2014;23(4-5):s505–12.

37. Kean TJ, Lin P, Caplan AI, Dennis JE. MSCs: Delivery Routes and Engraftment, Cell-Targeting Strategies, and Immune Modulation. Stem Cells Int. 2013;2013:732742.

38. Montarras D, Morgan J, Collins C, Relaix F, Zaffran S, Cumano A, et al. Direct isolation of satellite cells for skeletal muscle regeneration. Science. 2005;309(309):2064–7.

39. Bentivegna A, Roversi G, Riva G, Paoletta L, Redaelli S, Miloso M, et al. The Effect of Culture on Human Bone Marrow Mesenchymal Stem Cells: Focus on DNA Methylation Profiles. Stem Cells Int. 2016;2016:5656701.

40. Kirkland JL, Tchkonia T. Cellular Senescence: A Translational Perspective. EBioMedicine. 2017;21:21–8.

41. Turinetto V, Vitale E, Giachino C. Senescence in Human Mesenchymal Stem Cells: Functional Changes and Implications in Stem Cell-Based Therapy. International Journal of Molecular Sciences. 2016;17(17):1164-.

42. Schellenberg A, Stiehl T, Horn P, Joussen S, Pallua N, Ho AD, et al. Population dynamics of mesenchymal stromal cells during culture expansion. Cytotherapy. 2012;14(14):401–11.

43. Wagner W, Horn P, Castoldi M, Diehlmann A, Bork S, Saffrich R, et al. Replicative Senescence of Mesenchymal Stem Cells: A Continuous and Organized Process. PLoS ONE. 2008;3(3):e2213–e.

44. Hladik D, Höfig I, Oestreicher U, Beckers J, Matjanovski M, Bao X, et al. Long-term culture of mesenchymal stem cells impairs ATM-dependent recognition of DNA breaks and increases genetic instability. Stem Cell Research & Therapy. 2019;10(10):218-.

45. Musi N, Valentine JM, Sickora KR, Baeuerle E, Thompson CS, Shen Q, et al. Tau protein aggregation is associated with cellular senescence in the brain. Aging Cell. 2018;17(17):e12840.

46. Justice JN, Gregory H, Tchkonia T, LeBrasseur NK, Kirkland JL, Kritchevsky SB, et al. Cellular Senescence Biomarker p16INK4a+ Cell Burden in Thigh Adipose is Associated With Poor Physical Function in Older Women. J Gerontol A Biol Sci Med Sci. 2018;73(73):939–45.

47. Justice JN, Nambiar AM, Tchkonia T, LeBrasseur NK, Pascual R, Hashmi SK, et al. Senolytics in idiopathic pulmonary fibrosis: Results from a first-in-human, open-label, pilot study. EBioMedicine. 2019;40:554–63.

48. Docherty MH, O’Sullivan ED, Bonventre JV, Ferenbach DA. Cellular Senescence in the Kidney. J Am Soc Nephrol. 2019;30(30):726–36.

49. Roos CM, Zhang B, Palmer AK, Ogrodnik MB, Pirtskhalava T, Thalji NM, et al. Chronic senolytic treatment alleviates established vasomotor dysfunction in aged or atherosclerotic mice. Aging cell. 2016;15(15):973–7.

50. Zhu Y, Tchkonia T, Fuhrmann-Stroissnigg H, Dai HM, Ling YY, Stout MB, et al. Identification of a novel senolytic agent, navitoclax, targeting the Bcl-2 family of anti-apoptotic factors. Aging cell. 2016;15(15):428–35.

51. Khan N, Syed DN, Ahmad N, Mukhtar H. Fisetin: A Dietary Antioxidant for Health Promotion. Antioxidants & Redox Signaling. 2013;19(19):151–62.

52. Alleviation by Fisetin of Frailty, Inflammation, and Related Measures in Older Adults. clinicaltrialsgov 2021.

